# Benchmarking a highly selective USP30 inhibitor for enhancement of mitophagy and pexophagy

**DOI:** 10.1101/2021.04.28.441730

**Authors:** Emma V. Rusilowicz-Jones, Francesco G. Barone, Fernanda Martins Lopes, Elizabeth Stephen, Heather Mortiboys, Sylvie Urbé, Michael J. Clague

## Abstract

The deubiquitylase USP30 is an actionable target considered for treatment of conditions associated with defects in the PINK1/Parkin pathway leading to mitophagy. These include Parkinson’s disease and pulmonary fibrosis. We provide a detailed cell biological characterisation of a benzenesulphonamide molecule, compound 39, that has previously been reported to inhibit USP30 in an *in vitro* enzymatic assay. The current compound offers increased selectivity over previously described inhibitors. It enhances mitophagy and generates a signature response for USP30 inhibition following mitochondrial depolarisation. This includes enhancement of TOM20 and SYNJ2BP ubiquitylation and phosphoubiquitin accumulation, alongside increased mitophagy. In dopaminergic neurons, generated from Parkinson’s disease patients carrying loss of function Parkin mutations, compound 39 could significantly restore mitophagy to a level approaching control values. USP30 is located on both mitochondria and peroxisomes and has also been linked to the PINK1 independent pexophagy pathway. Using a fluorescence reporter of pexophagy expressed in U20S cells, we observe increased pexophagy upon application of compound 39 that recapitulates the previously described effect for USP30 depletion. This provides the first pharmacological intervention with a synthetic molecule to enhance peroxisome turnover.

## Introduction

The ubiquitin specific protease (USP) family of proteins represents an emerging focus for drug discovery efforts [1]. Amongst this family, USP30 is unique in exclusively localising to mitochondria and peroxisomes, by virtue of its transmembrane domain [2–4]. At mitochondria, it is optimally positioned to oppose the PINK1/Parkin mediated cascade of ubiquitylation that follows mitochondrial depolarisation and leads to mitophagy [5, 6]. Thus inhibition of USP30 represents an actionable target to correct pathologies associated with PINK1 or Parkin defects, such as Parkinson’s disease or pulmonary fibrosis [7, 8]. We and others have shown that USP30 localises to peroxisomes and that its depletion can lead to elevation of pexophagy without any effect on bulk autophagy [4, 9]. Peroxisome levels determine the abundance of ether linked lipids (plasmalogens) which are important for ferroptotic cell death in multiple cancer cells and are reduced in the brains of Alzheimer’s patients [10, 11]. Current tools to manipulate peroxisome turnover are limited to manipulation of lipids in the specific context of hepatocytes or crude approaches which stimulate general autophagy [12–14].

Recent studies have reported highly related cyanopyrrolidine USP30 covalent inhibitors that recapitulate signature effects of USP30 depletion in the context of acute mitochondrial depolarisation [15–17]. These include enhanced TOM20 ubiquitylation, increased mitophagy and greater accumulation of phosphoubiquitin [6, 15–18]. In the case of FT385 the specificity of the drug across a panel of DUBs was high at concentrations up to 200nM but it becomes less specific at higher concentrations. In the optimal concentration range it is recommended to define target engagement for each cell line under study using a competition assay with reactive ubiquitin probes [16, 19]. Phu et al. used higher concentrations of a related compound in a large scale proteomic analysis. By comparison with USP30^-/-^ cells, they were able to characterise both on and off target effects of the drug [15].

In light of these limitations, we decided to characterise a benzosulphonamide, from a series of compounds, described in the literature as specific USP30 inhibitors. Compound 39 (CMPD-39) has a reported IC_50_ of ~20nM in an *in vitro* assay of enzyme activity. In the same study a related compound was shown to accelerate the turnover of mitochondrial DNA in terminally differentiated C2C12 cells, but otherwise their biological effects are uncharacterised [20, 21]. Using USP30^-/-^ cells for comparison and off target assessment, our results validate a highly selective inhibition of USP30, which recapitulates the signature associated with structurally unrelated inhibitors. Moreover, we show for the first time that chemical inhibition of USP30 can increase basal pexophagy, to an extent consistent with previous observations of USP30 deletion and depletion [4, 9].

## Results

Previous work has reported IC_50_ values for a series of benzosulphonamide inhibitors, of which all those < 1 *μ*M were screened for selectivity, in comparison to three other USP family members [20, 21]. We have selected one of these compounds (CMPD-39, Figure 1A) for a more rigorous test of selectivity provided by the Ubiquigent DUB profiler screening platform comprised of >40 DUBs. CMPD-39 is a highly selective inhibitor of USP30 over two orders of magnitude of concentration (1-100*μ*M, Figure 1B). To test engagement of the inhibitor with USP30 once applied to cells, we assayed competition with Ub-propargylamide (Ub-PA). Ub-PA covalently binds to the USP30 active site leading to a characteristic upshift on an SDS-PAGE gel [22]. If a drug blocks access to the catalytic site then the probe modification is blocked and the apparent molecular weight of USP30 shifts down accordingly. Applying CMPD-39 to SHSY5Y neuroblastoma cells shows strong competition for Ub-PA in the sub *μ*M range of concentrations (Figure 1C). A robust proxy read-out for target engagement is the enhancement of TOM20 ubiquitylation following mitochondrial depolarisation [6, 16]. This second assay indicates a maximal effect with 200 nM CMPD-39 applied to RPE1-YFP-Parkin cells cells (Figure 1D).

**Figure 1.**
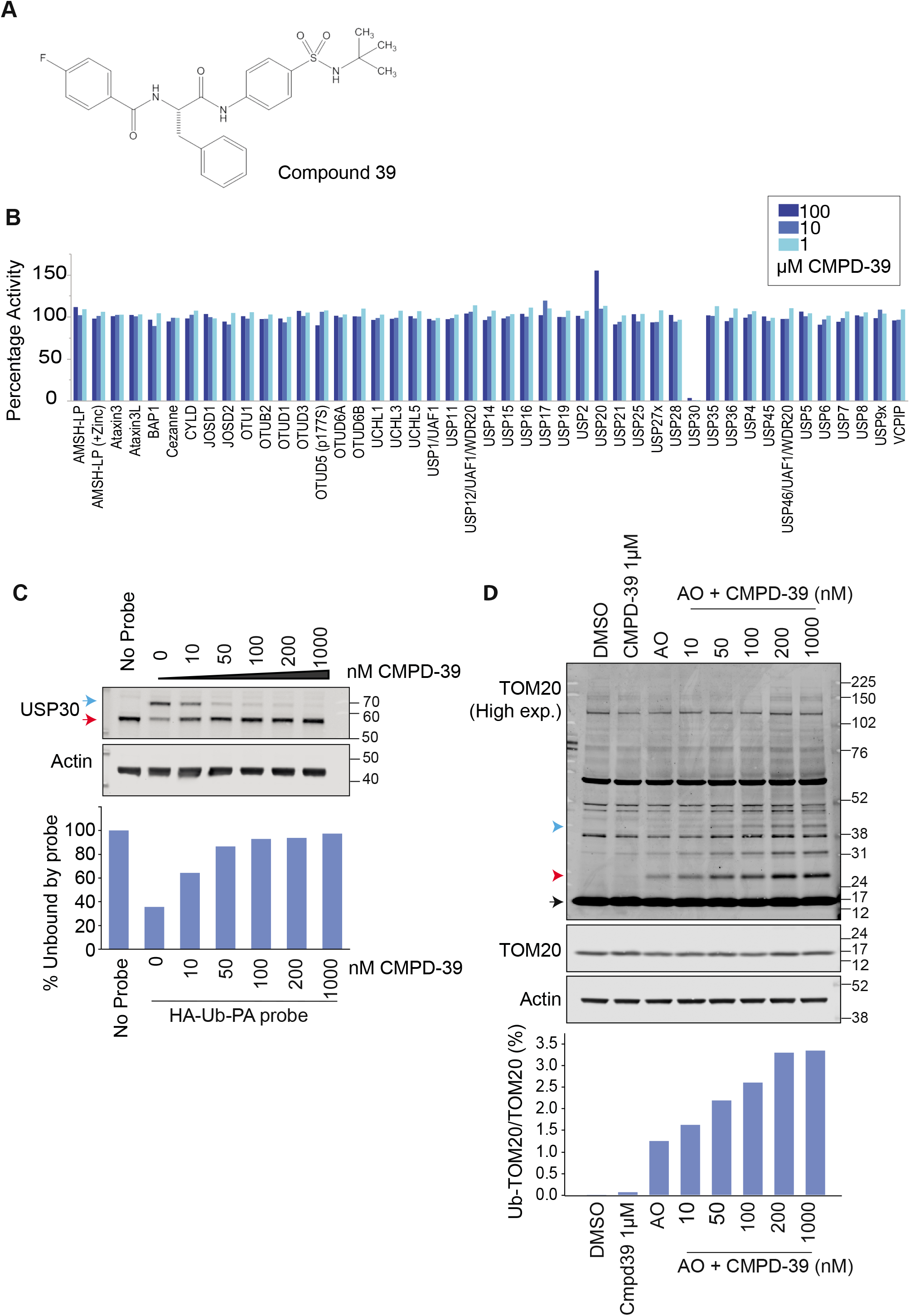
Compound 39 is a selective USP30 inhibitor. (A) Chemical structure of CMPD-39. (B) DUB specificity screen (DUB profiler, Ubiquigent) with 1-100*μ*M CMPD-39. (C) Activity based ubiquitin probe assay shows that CMPD-39 engages USP30 in cells at nanomolar concentrations in intact SHSY5Y cells. Samples were incubated with CMPD-39 for 2 hours at the indicated concentrations, then incubated with HA-Ub-PA probe for 10 minutes at 37°C and immunoblotted as shown. Red arrow indicates unbound USP30, Blue arrow represents probe bound USP30. (D) Inhibition of USP30 enhances the ubiquitylation of TOM20 in YFP-Parkin over-expressing hTERT-RPE1 cells in a concentration dependent manner in response to mitophagy induction. Cells were treated for 1 hour with DMSO or Antimycin A and Oligomycin A (AO; 1 *μ*M each) in the absence or presence of CMPD-39 at the indicated concentrations, lysed and analysed by western blotting. Black arrow indicates unmodified TOM20, ubiquitylated species are indicated by red (mono-ubiquitylated) or blue arrow heads. Quantitation shows percentage mono-ubiquitylated.

We next took advantage of previously described isogenic USP30^-/-^ RPE1-YFP-Parkin and SHSY5Y cell lines to demonstrate the target dependence of three signature effects of USP30 inhibition. As previously indicated, depolarisation (A/O) induced TOM20 ubiquitylation is enhanced by 200 nM CMPD-39. However in USP30^-/-^ cells the signal is already maximally elevated and is impervious to the inhibitor (Figure 2A,B,D,E). The same pattern holds for a second biomarker, SYNJ2BP, identified by global ubiquitomic analysis of the response to FT385, a cyanopyrrolidine class of USP30 inhibitor (Figure 2A,C,D) [16]. Mitochondrial depolarisation leads to the accumulation of the kinase PINK1 which phosphorylates ubiquitin (pUb) and Parkin at Ser65 [23, 24]. This serves to recruit and activate Parkin, setting off a cascade of ubiquitylation at the mitochondrial surface [25]. USP30 inhibition with cyanopyrrolidine inhibitors has the effect of modestly enhancing the PINK1 dependent accumulation of pUb on mitochondria [16, 17]. In SHSY5Y cells this is most apparent between the 38 and 76kD molecular weight range and is now reproduced with CMPD-39 or knock-out of USP30 with no additive effect (Figure 2D,F).

**Figure 2.**
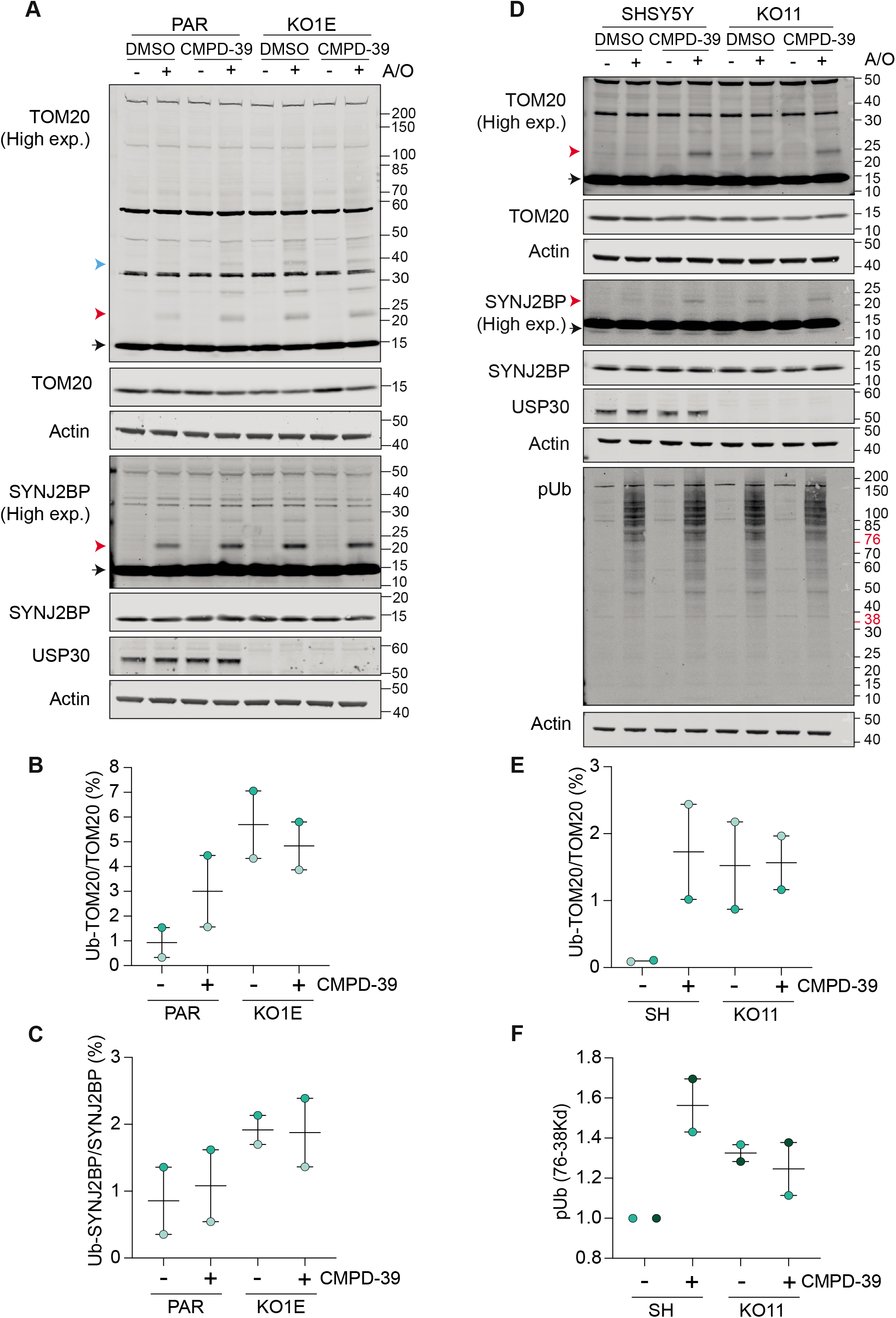
CMPD-39 promotes depolarisation dependent ubiquitylation of previously described USP30 substrates. (A) USP30 inhibitor (CMPD-39) treatment of parental hTERT-RPE1-YFP Parkin cells phenocopies USP30 deletion (KO1E) by promoting TOM20 and SYNJ2BP ubiquitylation. TOM20 and SYNJ2BP ubiquitylation is unaffected by CMPD-39 in the USP30 KO (KO1E) cells. Cells were treated for 1 hour with or without AO (1 *μ*M) in the absence or presence of 200 nM CMPD-39. (B,C) Graphs show quantification of mono-ubiquitylated TOM20 and SYNJ2BP in AO-treated samples as a percentage of total for two independent experiments (error bars indicate the range). (D) USP30 inhibitor (CMPD-39) treatment of cells expressing endogenous Parkin (SHSY5Y cells) similarly phenocopies USP30 deletion (KO11) by promoting TOM20 and SYNJ2BP ubiquitylation, as well as increasing levels of phospho-Ser65 Ubiquitin (pUb). Cells were treated for 4 hours with or without AO (1 *μ*M) in the absence or presence of 1 *μ*M CMPD-39. (E,F) Graphs show quantification of mono-ubiquitylated TOM20 as a percentage of total TOM20 (E) and the pUb signal (F) in the 38–76 kD range, for two independent experiments represented by (D) (error bars indicate the range). (A, D): Black arrow indicates unmodified species, and red and blue arrowheads indicate the mono- and multi-ubiquitylated species respectively.

So far, the biochemical signatures we have presented require acute mitochondrial depolarisation, which represents a non-physiological condition. Under basal conditions only a small fraction of mitochondria is undergoing mitophagy. We have quantitated this fraction using SHSY5Y cells stably expressing the fluorescent mitophagy reporter mitoQC [26]. At 1 *μ*M CMPD-39 the number and size of mitolysosomes is increased, in line with previous observations using FT385 (Figure 3A-C) [16]. Interest in the therapeutic potential of USP30 inhibitors has been driven by their application in Parkinson’s disease, for which some patients have loss of function mutations in the Parkin (PARK2) gene. We next turned to dopaminergic neurons generated from patient derived fibroblasts, via induced neuronal progenitor cells [27]. In this experiment we quantitate the co-localisation of lysotracker with the mitochondrial marker tetramethylrhodamine as an indicator of mitophagy in conjunction with the Opera Phenix screening platform, as previously described [27]. The level of mitophagy in neurons from two Parkin compound heterozygotic mutant patients was reduced compared with two controls, in line with previous observations [27]. Upon application of CMPD-39 to the neuronal cultures for 24 hours, the control samples showed a trend towards increased mitophagy. Most notably both Parkin mutant cell lines showed a statistically significant increase, such that their mitophagy index was restored to near control levels (Figure 3D).

**Figure 3.**
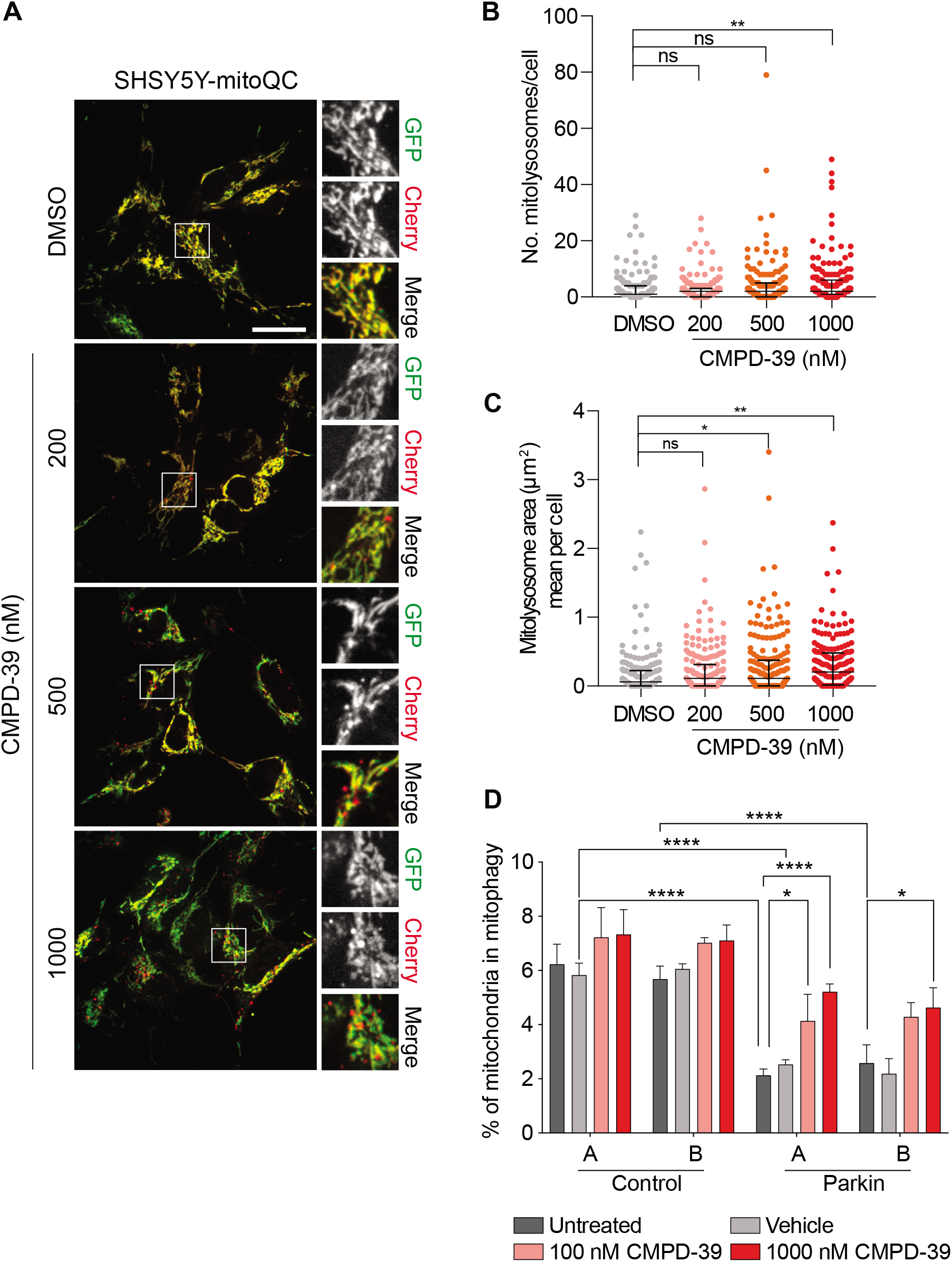
Enhancement of basal mitophagy by pharmacological inhibition of USP30. (A) Representative images of SHSY5Y-mitoQC cells stably expressing mitoQC. Cells were treated with DMSO or CMPD-39 (200, 500 or 1000 nM) for 96 hours before imaging. Scale bar 20 *μ*M. (B-C) Quantification for the data from three independent experiments is shown. Error bars indicate standard deviation; 65 cells were quantified per experiment. (B) Graph illustrates the number of mitolysosomes. **P<0.01, Oneway ANOVA with Dunnett’s multiple comparisons test. (C) Graph shows the mean mitolysosome area per cell. *P<0.05, **P<0.01, One-way ANOVA with Dunnett’s multiple comparisons test, (D) mitophagy index for patient derived dopaminergic iNeurons derived from two control individuals and two individuals with Parkin loss of function mutations.

Finally we examined the effect on basal pexophagy, which is a PINK/Parkin independent pathway, but can nevertheless be enhanced by USP30 depletion without a corresponding gain in non-selective autophagy [4, 9]. For this, we used U2OS cells stably transfected with Keima-SKL, a fluorescence reporter for pexophagy [4]. When peroxisomes reach the acidic lysosome compartment, the excitation spectrum of the Keima fluorophore undergoes a shift that is then pseudo-coloured as red in the illustrative images. CMPD-39 induced a strong increase in basal pexophagy (Figure 4), in line with our previous observations, obtained by USP30 depletion with specific siRNAs [4]. To our knowledge, this is the first example of a synthetic compound with a clearly defined target, that can promote pexophagy.

**Figure 4.**
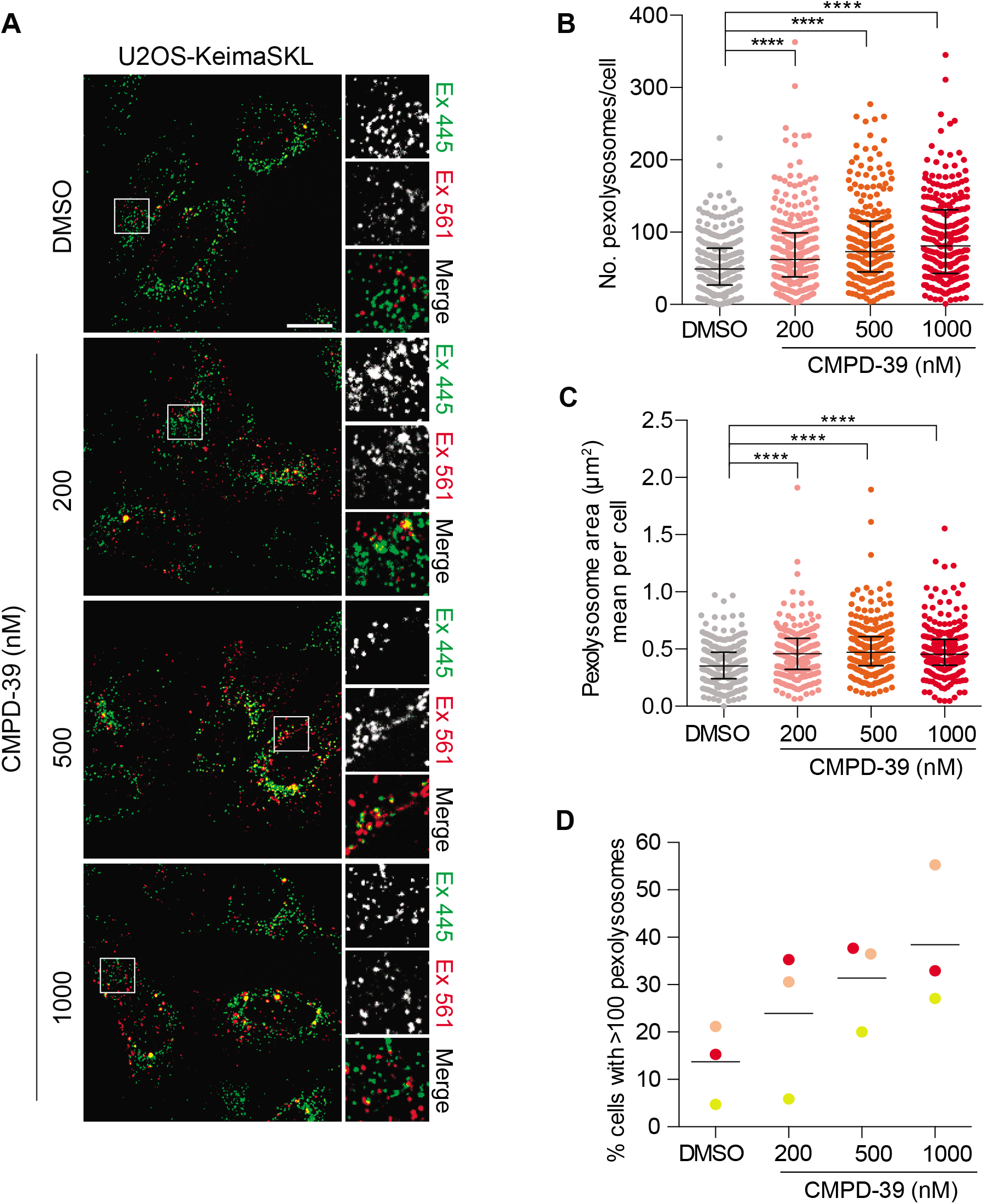
Enhancement of basal pexophagy by pharmacological inhibition of USP30. (A) Representative images of U2OS-KeimaSKL cells stably expressing KeimaSKL. Cells were treated with DMSO or CMPD-39 (200, 500 or 1000 nM) for 96 hours before imaging. Scale bar 20um. Quantification of the data is derived from three independent experiments. Error bars indicate standard deviation; 85 cells were quantified per experiment. (B) Graph indicates the number of pexolysosomes in U2OS-KeimaSKL cells. One-way ANOVA with Dunnett’s multiple comparisons test, ****P<0.0001. (C) Graph indicates the mean area of pexolysosomes per cell. One-way ANOVA with Dunnett’s multiple comparisons test, ****P<0.0001. (D) Percentage of U2OS-KeimaSKL cells with 100 or more pexolysosomes per cell. Different colours represent data from three independent experiments.

## Discussion

Across the neurodegeneration landscape actionable targets are still relatively scarce. Much attention has focused on enhancing mitophagy. In Parkinson’s disease this has been driven by the understanding that two Parkinson’s associated genes PINK1 and Parkin combine to promote mitophagy and that this pathway is deficient in patients bearing loss of function mutations [28]. Defects in mitochondria are also evident in idiopathic patients or those with mutations in other PD associated genes such as LRRK2 [29–31]. Furthermore, mitophagy is impaired in Alzheimer’s Disease [32, 33] and loss of function mutations in the mitophagy regulator TBK1 and the adaptor protein, optineurin, are associated with Amyotrophic Lateral Sclerosis (ALS) [34].

The PINK1/Parkin pathway is the best understood mitophagy pathway and is propelled by a Parkin-mediated cascade of ubiquitylation at the mitochondrial surface. As the most prominent mitochondrial deubiquitylase, USP30 is ideally placed to limit this pathway [3]. If that is the case, then chemically inhibiting USP30 offers a therapeutic opportunity to enhance mitophagy. A further attractive feature of USP30 inhibition is that it is a non-essential gene, and this is a necessary condition for any long-term treatment option. The best characterised USP30 inhibitors to date are a series of cyanopyrrolidines, for which a consensus biochemical and cell physiological signature is beginning to emerge [5, 16, 17]. One limitation of the use of these compounds is that they become less specific at higher concentrations and this can become problematic if the cellular uptake and target engagement efficiency is unknown [15, 17]. We wondered if the recently published series of benzosulphonamide USP30 inhibitors might represent a structurally independent alternative, offering a greater dynamic range of concentration over which inhibition is achieved and specificity is retained [20]. The cell physiological characterisation of these compounds has hitherto been rudimentary. We now show that CMPD-39 is a highly selective USP30 inhibitor, that can achieve maximal USP30 inhibition at <200 nM in SHSY5Y cells but retains selectivity amongst DUB family members even up to 100 *μ*M. Our data obtained with this molecule suggest that it is likely to provide a more robust tool compound than the cyanopyrrolidines. We reinforce the consensus signature for USP30 inhibition i.e. (i) increased TOM20 and SYNJ2BP ubiquitylation following mitochondrial depolarisation, (ii) moderately enhanced pUb accumulation following mitochondrial depolarisation, (iii) enhancement of basal mitophagy.

It is known that dopaminergic neurons derived from patients carrying loss of function Parkinson’s mutations show lower levels of mitophagy [27]. Here we now show that this defect can be restored by a specific inhibitor of USP30, providing further encouragement for pre-clinical development of these compounds. Similar results have recently been reported in pre-print form for a set of cyanopyrrolidine inhibitors [17].

Peroxisomes are strongly linked to neurodegenerative disease [35, 36]. They support oxidation of various fatty acids and regulate redox conditions. They are involved in synthesis of ether linked phospholipids, notably plasmalogens, that are highly enriched in the nervous system. A fraction of USP30 associates with peroxisomes and its loss leads to an increase in pexophagy [4, 9, 37]. siRNA depletion or knock-out of USP30 in RPE1 cells enhances pexophagy and importantly this effect can be rescued by re-expression of USP30, but not a catalytically inactive mutant [4]. We now show that chemical inhibition of USP30 also enhances pexophagy in U2OS cells. The ability to acutely manipulate this process in a specific manner has been lacking and we anticipate that CMPD-39 could become a widely adopted tool compound for this alternative application. In summary, benzosulphonamide USP30 inhibitors, specifically CMPD-39, offer an important new class of tool compound for enhancement of mitophagy and pexophagy.

## Acknowledgements

This study was funded by Celgene and Bristol Myers Squibb. We thank Richard Hargreaves and Jeff Schkeryantz for their encouragement and enablement of this work. RPE1-Parkin-YFP cells were a kind gift of Jon Lane (University of Bristol). FB is a Wellcome Trust funded PhD student.

## Methods

### Cell culture

hTERT-RPE1-YFP-PARKIN, SHSY5Y, SHSY5Y-mitoQC (mCherry-GFP-Fis1(101-152)) and U2OS-Keima-SKL were routinely cultured in Dulbecco’s Modified Eagle’s medium DMEM/F12 supplemented with 10% FBS and 1% non-essential amino acids. USP30^-/-^ cells were generated as described in Rusilowicz-Jones et al. [16]. Primary fibroblasts were obtained from Coriell Cell Repository (coriell.org); control A GM13335 (M57), control B ND29510 (F55), Parkin A ND30171 (M54, Arg42Pro, EX3Del), Parkin B ND40067 (F44, EX4-7DEL c203_204 DEL AG). Fibroblasts were cultured in EMEM as previously described [38]. iNPC’s were generated as previously described [39]. iNPC’s were maintained in DMEM/Ham F12 (Invitrogen); N2, B27 supplements (Invitrogen) and FGFb (Peprotech) in fibronectin (Millipore) coated tissue culture dishes and routinely sub-cultured every 2–3 days using accutase (Sigma) for detachment. Neurons were differentiated from iNPC’s as previously described [40]. Briefly, iNPCs are plated in a 6-well plate and cultured for 2 days in DMEM/F-12 medium with Glutamax supplemented with 1% NEAA, 2% B27 (Gibco) and 2.5 *μ*M of DAPT (Tocris). On day 3, DAPT is removed, and the medium is supplemented with 1 *μ*M smoothened agonist (SAG; Millipore) and FGF8 (75 ng/ml; Peprotech) for additional 10 days. Neurons are replated and subsequently SAG and FGF8 are withdrawn and replaced with BDNF (30 ng/ml; Peprotech), GDNF (30 ng/ml; Peprotech), TGF-b3 (2 mM; Peprotech) and dcAMP (2 mM, Sigma) for 15 days.

### Antibodies and reagents

Antibodies and other reagents used were as follows: anti-USP30 (Sigma HPA016952, 1:500), anti-TOM20 (ProteinTech 11802-1-AP, 1:1,000), mouse anti-actin (ProteinTech 66009-1-Ig, 1:10,000) anti-phospho-Ubiquitin Ser65 (Millipore, ABS1513-I, 1:1000), anti-SYNJ2BP (Sigma HPA000866, 1:1000), oligomycin A (SIGMA 75351), antimycin A (SIGMA A8674).

### Preparation cell lysates and Western blot analysis

Cultured cells were lysed with NP-40 (0.5% NP-40, 25 mM Tris–HCl pH 7.5, 100 mM NaCl, 50 mM NaF) lysis buffer and routinely supplemented with mammalian protease inhibitor (MPI) cocktail (Sigma) and Phostop (Roche). Proteins were resolved using SDS–PAGE (Invitrogen NuPage gel 4–12%), transferred to nitrocellulose membrane, blocked in 5% milk or 5% BSA in TBS supplemented with Tween-20, and probed with primary antibodies overnight. Visualisation and quantification of Western blots were performed using IRdye 800CW and 680LT coupled secondary antibodies and an Odyssey infrared scanner (LI-COR Biosciences, Lincoln, NE).

### Activity probe assay

Cells were mechanically homogenised in HIM buffer (200 mM mannitol, 70 mM sucrose, 1 mM EGTA, 10 mM HEPES-NaOH, pH 7.4) supplemented with 1 mM DTT to obtain post-nuclear supernatants (PNS). Briefly SHSY5Y cells were washed with ice-cold PBS (+ 1mM DTT) and then collected by scraping and centrifugation at 1,000 g for 2 minutes. Cell pellets were washed with HIM buffer and then resuspended in HIM buffer (+DTT). Cells were mechanically disrupted by shearing through a syringe with a 27G needle, followed by 3 times though an 8.02 mm diameter “cell cracker” homogeniser using a 8.01 mm diameter ball bearing. The resulting homogenate was cleared from nuclei and unbroken cells by centrifugation at 600 g for 10 minutes to obtain the PNS. Homogenates were incubated with HA-Ub-PA probe at 1:100 (w/w) for 10 minutes at 37°C. The reaction was stopped by the addition of sample buffer and heating at 95°C. To test drug engagement with USP30, intact cells were treated with CMPD-39 for 2 hours at 37°C prior to homogenisation followed by probe incubation.

### Live cell imaging

SHSY5Y cells stably expressing mCherry-GFP-Fis1(101-152) (SHSY5Y mitoQC) or U2OS cells stably expressing Keima-SKL were treated every 24 hrs over a 96 hrs timecourse with 200, 500 nM and 1*μ*M of CMPD-39. Cells were replated onto an IBIDI μ-Dish (2×105) two days before live-cell imaging with a 3i Marianas spinning disk confocal microscope (63x oil objective, NA 1.4, Photometrics Evolve EMCCD camera, Slide Book 3i v3.0). Cells were randomly selected using the GFP (mitoQC) or Keima-445 (KeimaSKL) signal and images acquired sequentially using the following settings. SHSY5Y mitoQC: 488 nm laser, 525/30 emission; 561 nm laser, 617/73 emission, U2OS-KeimaSKL: 445 nm laser, 525/30 emission; 561 nm laser, 617/73 emission.

In order to assess mitophagy in live iNeurons a modified version of the protocol in Schwartzentruber et al, 2020 was adopted [27]. Briefly, cells were incubated for one hour at 37°C with 80 nM tetramethylrhodamine (TMRM), 1 *μ*M LysoTracker Deep Red (Invitrogen), 1 *μ*M MitoTracker Green (Invitrogen) and 1 *μ*M Hoechst, before washing to remove fluorescent probes. Images were captured by the Opera Phenix system (Perkin Elmer) in time lapse, every 18 minutes in the same fields of view, minimum 8 fields of view per well and 5 z slices. Images were taken using the following settings, 488nm laser, emission 500-550; 568nm laser, emission 570-630; 647nm laser, emission 650-760; 405nm laser, emission 435-480.

### Image quantification

Analysis of mitophagy and pexophagy levels in SHSY5Y-mitoQC and U2OS-Keima-SKL cells was performed using the ‘mito-QC Counter’ implemented in FIJI v2.0 software (ImageJ, NIH) as previously described [41]. For mitophagy the following parameters were used: Radius for smoothing images = 1.5, Ratio threshold = 0.8, and Red channel threshold = mean. For pexophagy the following parameters were used: Radius for smoothing images = 1.5, Ratio threshold = 1.9, and Red channel threshold = mean. Mitophagy and pexophagy analysis was performed for three independent experiments with 65 and 85 cells per condition respectively. One-Way ANOVAs with Dunnett’s multiple comparisons were performed using GraphPad Prism 9. P-values are represented as *P<0.05, **P<0.01, ****P<0.0001. Images generated from the live iNeuron imaging experiments were analysed using Harmony (Perkin Elmer software). We developed protocols in order to segment nucleus, image regions containing cytoplasm, mitochondria, lysosomes, autolysosomes containing mitochondria. Maximal projection images were used for analysis. Mitochondria contained within lysosomes segmentation was set up in such a way to identify a mitophagy event when the overlap between mitochondria and lysosome was 100%.

